# Different Coupling Modes Mediate Cortical Cross-Frequency Interactions

**DOI:** 10.1101/016212

**Authors:** Randolph F. Helfrich, Christoph S. Herrmann, Andreas K. Engel, Till R. Schneider

## Abstract

Cross-frequency coupling (CFC) has been suggested to constitute a highly flexible mechanism for cortical information gating and processing, giving rise to conscious perception and various higher cognitive functions in humans. In particular, it might provide an elegant tool for information integration across several spatiotemporal scales within nested or coupled neuronal networks. However, it is currently unknown whether low frequency (theta/alpha) or high frequency gamma oscillations orchestrate cross-frequency interactions, raising the question of who is master and who is slave. While correlative evidence suggested that at least two distinct CFC modes exist, namely phase-amplitude-coupling (PAC) and amplitude-envelope-correlations (AEC), it is currently unknown whether they subserve distinct cortical functions. Novel non-invasive brain stimulation tools, such as transcranial alternating current stimulation (tACS), now provide the unique opportunity to selectively entrain the low or high frequency component and study subsequent effects on CFC. Here, we demonstrate the differential modulation of CFC during selective entrainment of alpha or gamma oscillations. Our results reveal that entrainment of the low frequency component increased PAC, where gamma power became preferentially locked to the trough of the alpha oscillation, while gamma-band entrainment reduced alpha power through enhanced AECs. These results provide causal evidence for the functional role of coupled alpha and gamma oscillations for visual processing.

## Introduction

Cognition and conscious perception are thought to arise from neuronal interactions between functionally specialized but widely distributed cortical regions (Siegel et al., 2012). While phase synchronization between task-relevant cortical areas might integrate information across several spatial scales (Fries, 2005), cross-frequency coupling (CFC) has been suggested to constitute a flexible mechanism for information integration across temporal scales (Canolty and Knight, 2010). Hence, it might serve as a key mechanism for selective gating and processing of information within coupled or nested cortical networks and thus, subserve numerous cognitive functions in humans (Canolty and Knight, 2010; Voytek et al., 2010; Engel et al., 2013). However, the functional role of CFC is currently extensively under debate, given that (I) various methodological constrains hamper its interpretation (Aru et al., (II) the evidence supporting its role for cognitive processing was only correlative in nature (Canolty and Knight, 2010; Voytek et al., 2010) and (III) it remained unclear whether different coupling modes (e.g. PAC or AEC) subserve distinct cortical functions. In particular, alpha-gamma PAC has been suggested to constitute a powerful mechanism to organize visual processing (Spaak et al., 2012; Jensen et al., 2014), while theta-gamma PAC might subserve memory processes and long-range cortico-cortical communication (Tort et al., 2009; Axmacher et al., 2010; Canolty and Knight, 2010; Lisman and Jensen, 2013). Most studies focused on PAC (Canolty and Knight, 2010; Voytek et al., 2010), where gamma power is preferentially phase-locked to the trough of the theta (4–7 Hz) or the alpha (8-12 Hz) rhythm. However, it has recently been observed that the cerebral cortex exhibits a large-scale correlation structure which is independent from phase-locked signaling and can best be analyzed by quantifying AEC (Hipp et al., 2012; Engel et al., 2013). While first applied to uncover envelope correlations within one frequency range, AEC can also be applied to the cross-frequency domain (Helfrich et al., 2014a). Figure 1A depicts a schematic how both CFC measures (PAC and AEC) were obtained from raw data.

**Figure 1.**
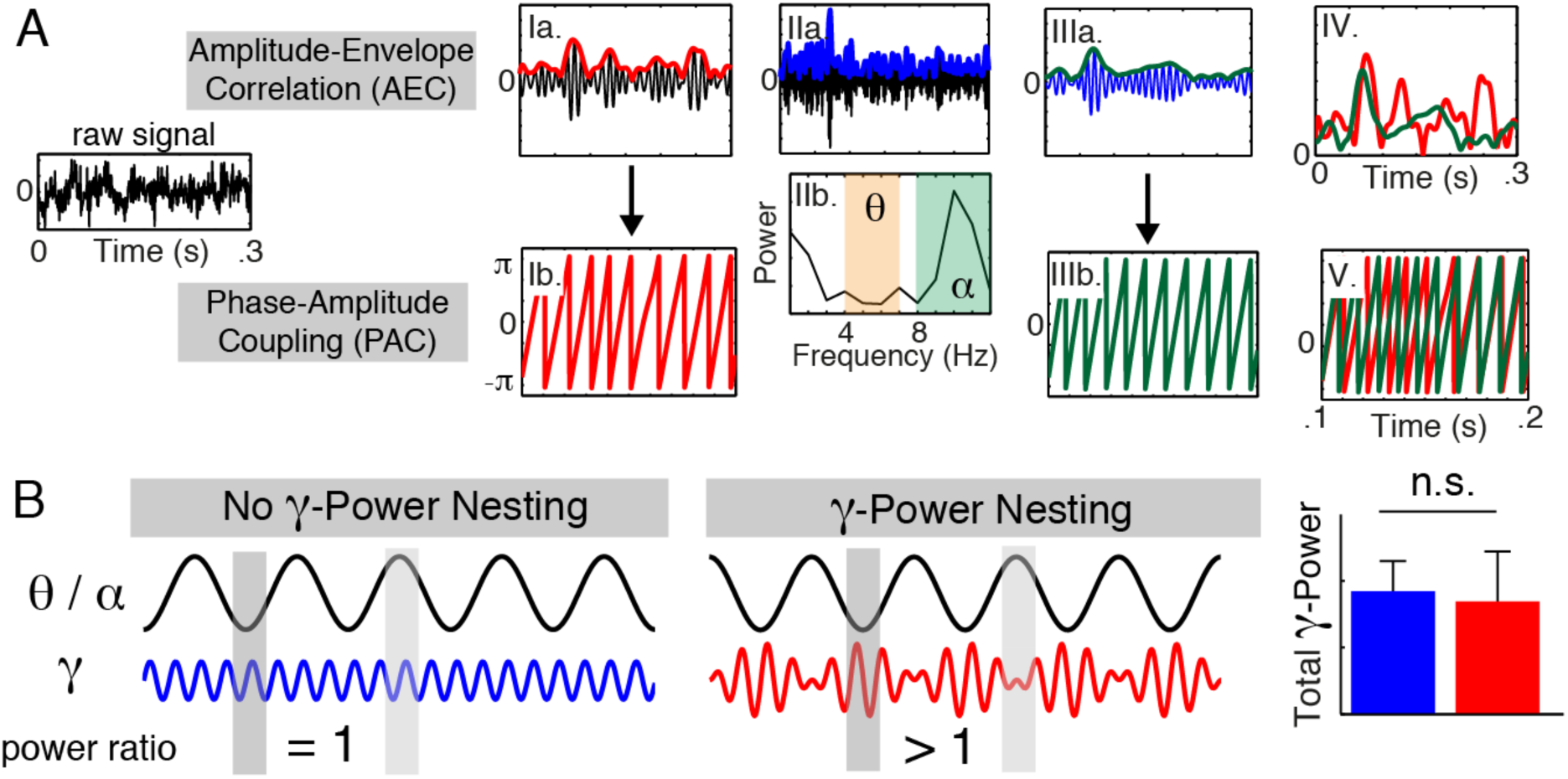
Schematic illustration of CFC analysis. (A) Ia. Low frequency band-pass filtered signal (black) and the corresponding envelope (red) and instantaneous phase (Ib.). IIa. Gamma band-pass filtered signal (black) and the corresponding envelope (blue). IIb. Fast Fourier Transform (FFT) of the gamma envelope indicates that lower frequency components modulate the instantaneous gamma amplitude. Shaded areas highlight theta (orange, 4–7 Hz) and alpha (green, 8–12 Hz) frequency ranges that were utilized for subsequent analyses. IIIa. The gamma envelope (blue) was filtered in the low frequency range and then its envelope (green) and phase (IIIb.) were extracted. IV. Superposition of low frequency amplitude (Ia.) and gamma envelope (IIIa.), which were then used for AEC analysis by means of Pearson’s linear correlation coefficient (Fisher-Z-transformed). V. Superposition of low frequency phase (Ib.) and gamma phase after band-pass filtering (IIIb.). Their interaction was analyzed by means of the phase-locking value (PLV; Voytek et al., 2010) to obtain the PAC. (B) Schematic of the relationship of the low frequency (theta/alpha) and the gamma oscillation to illustrate the nesting effect, i.e. that the gamma amplitude is modulated by low frequency fluctuations. Note that absolute power values do not differ between conditions. However, in the case of nesting gamma power becomes preferentially locked to the trough of the low frequency oscillation. The power ratio was calculated as gamma power (trough-locked; shaded dark grey) divided by the gamma power (peak-locked; shaded light grey).

To investigate the mechanisms underlying the alpha-gamma interplay, we took advantage from a novel non-invasive brain stimulation technique, namely transcranial alternating current stimulation (tACS; Thut et al., 2011; Herrmann et al., 2013), which has recently been demonstrated to selectively entrain neuronal oscillations within canonical frequency boundaries (Ozen et al., 2010; Helfrich et al., 2014b). Thus, this approach provided the unique opportunity to selectively drive one spectral component and study subsequent cross-frequency effects in coupled frequency bands. Here, we reanalyzed data from two combined tACS-EEG (electroencephalography) studies on visual perception. Our aim was to study how cross-frequency interactions were modulated when either alpha (8–12 Hz; Helfrich et al., 2014b) or gamma (> 35 Hz; Helfrich et al., 2014a) oscillations were entrained by tACS, i.e., when one of these processes was set up as the master through synchronization to the external driving force.

## Material and Methods

### Participants

A total of 30 healthy, right-handed participants were recruited from the University of Oldenburg, Germany and the University Medical Center Hamburg-Eppendorf, Germany. Sixteen subjects (8 female; mean age: 24.6 ± 2.8 years; 2 excluded due to technical difficulties during data acquisition) participated in study 1 and fourteen subjects (7 female; mean age: 27.5 ± 6.7 years) participated in study 2. All participants reported no history of neurological or psychiatric disease and were medication-free during the experiments. They all had normal or corrected-to-normal vision and provided written informed consent according to the local ethics committee approval and the Declaration of Helsinki.

### General Procedure

All volunteers participated in one or two sessions of the experiments carried out within one week. After preparation of EEG and tACS electrodes, participants completed a training session to familiarize the volunteers with the visual stimuli to ensure adequate task performance. Participants reported their percepts by button presses with their right hand. Then all participants were familiarized with skin sensations and phosphenes, which have been reported in previous tACS studies (Kanai et al., 2010; Schutter and Hortensius, 2010). However, all participants indicated that the stimulation intensity was below skin sensation and phosphene threshold. In both studies the sham condition preceded active stimulation to avoid carry-over effects (Neuling et al., 2013).

### Data Acquisition

#### EEG recordings

All experiments were conducted with participants seated comfortably in a recliner in a dimly lit, electrically shielded room to avoid line noise interference. EEG electrodes were mounted in an elastic cap (Easycap, Herrsching, Germany) prepared with a slightly abrasive electrolyte gel (Abralyt 2000, Easycap, Herrsching, Germany). Ground-free EEG recordings (impedances < 20 kΩ, referenced to the nose tip) were obtained using BrainAmp amplifiers (Brain Products GmbH, Gilching, Germany) from 59 electrodes in study 1 and 31 electrodes in study 2. Signals were sampled at 1000 or 5000 Hz, amplified in the range of ± 16.384 mV at a resolution of 0.5 *μ*V and stored for offline analyses.

#### Electrical stimulation

In accordance with current safety limits (Nitsche et al., 2008), transcranial stimulation was applied by a battery-operated stimulator (DC-Stimulator Plus, NeuroConn, Ilmenau, Germany) via two rubber electrodes in study 1 (5x7 cm^2^, NeuroConn, Illmenau, Germany) and via ten Ag/AgCl electrodes in study 2 (12mm diameter, Easycap, Herrsching, Germany; resulting in a combined electrode area of approx. 11.3 cm^2^). Electrode placement was chosen in accordance with previous electrical brain stimulation studies targeting the extra-striate visual cortex (Neuling et al., 2013; Strüber et al., 2014). The combined impedance of all electrodes was kept below 5 kΩ, as measured by NeuroConn tACS device, using Ten20 conductive paste (Weaver and company, Aurora, Colorado) or Signa electrolyte gel (Parker Laboratories Inc., Fairfield, NJ, USA). A sinusoidally alternating current of 1000 *μ*A was applied at 10 or 40 Hz continuously for 20 minutes during each session. A tACS trigger was inserted every 30 cycles during the zero crossing of the external sine wave. During sham and real stimulation the current was ramped up to 1000 *μ*A, but discontinued after 20 seconds during the sham condition. All subjects confirmed that stimulation was acceptable and mainly noticeable during the ramp-in phase. It did not induce painful skin sensations or phosphenes. On debriefing, subjects indicated that they could not distinguish between sham and stimulation.

### Data Analysis

Data analysis was performed using MATLAB (The MathWorks Inc., Natick, MA, USA), EEGLAB (Delorme and Makeig, 2004); http://sccn.ucsd.edu/eeglab/), FieldTrip (Oostenveld et al., 2011); http://www.ru.nl/donders/fieldtrip/ and customized MatLab Code.

All preprocessing steps were performed with EEGLab after tACS artifact rejection (Helfrich et al., 2014a, 2014b). The recorded EEG was filtered using two-pass finite element response filters in the range from 1 - 100 Hz. Segments containing excessive noise, saccades or muscle artifacts were removed after visual inspection. Data were epoched into non-overlapping three-second segments and exported to Fieldtrip for further analysis, as described previously. Instantaneous power and phase of different frequency bands were obtained after band-pass filtering of the data and subsequent Hilbert transformation as described below.

Cross-frequency interactions between the slow oscillation (theta: 4 - 7 Hz; alpha: 8 – 12 Hz) and gamma (46 – 70 Hz) were assessed for every channel, condition and subject separately. The gamma-range was chosen in accordance with a recent report to keep results comparable (Helfrich et al., 2014a). In order to calculate the amplitude-envelope-correlation between both, we first band-pass filtered every trial in the low frequency range and in the gamma-range, similar to the approach, which had been introduced and validated before (Voytek et al., 2010).

Therefore, we utilized a two-way, zero phase-lag, finite impulse response (FIR) filter (eegfilt.m function in EEGLab toolbox; Delorme and Makeig, 2004) to prevent phase distortion. Hilbert’s transform was applied to the band-pass filtered low frequency signal to extract its amplitude and the instantaneous phase. Then a second Hilbert transform was utilized to extract the amplitude of the gamma oscillation, which was band-pass filtered in the low frequency range. Subsequently, a third Hilbert transform was used to extract the envelope and the instantaneous phase of the low-frequency band-pass filtered gamma amplitude. Amplitude-envelope-correlations (AEC) between the low frequency amplitude and the gamma envelope in the respective frequency range were calculated for every trial separately. Subsequently, Pearson’s linear correlation values were Fisher-Z-transformed and averaged. Phase-amplitude coupling (PAC) was calculated by means of the phase-locking technique (Lachaux et al., 1999; Voytek et al., 2010) between the instantaneous phase of the low-frequency oscillation and the instantaneous phase of gamma oscillation after band-pass filtering in the low-frequency range. Figure 1A provides a brief overview how both measures were obtained.

To calculate the gamma power relative to the low frequency oscillation, the analytic amplitude of the gamma-filtered signal was grouped into four equally sized phase-bins, relative to the phase of the low frequency oscillation (centered on the trough, the peak and the two zero-crossings). Then the amplitude values were squared to obtain the oscillatory power. In order to determine the amount of nesting of the gamma power relative to the low oscillatory phase, a power ratio (PR) was calculated as gamma power (locked to the trough of the slow frequency component) divided by the gamma power (locked to the peak of the slow frequency component). In case of no nesting of gamma power, the value is 1, i.e. the gamma power does not differ between peak and trough. Values > 1 indicate increased nesting, i.e. higher gamma power during the trough than during the peak of low frequency oscillation, while values < 1 point to the opposite. Figure 1B illustrates the nesting effect.

### Statistical analyses

We utilized Bonferroni-corrected t-tests according to the initial hypotheses at the channels-of-interest. Therefore, PAC and AEC analyses were corrected for 4 comparisons (p = 0.05/4 = 0.0125), while the power nesting analyses were corrected for 2 comparisons (p = 0.05/2 = 0.025). Effect sizes were quantified by means of Cohen’s d (Cohen, 1988), where values > 0.8 highlight a strong effect, values < 0.4 indicate no effect. Intermediate values signal a moderate effect size. All values reported are mean ± standard deviation. The regional specificity of the observed effects was confirmed with cluster-based permutation statistics as implemented in Fieldtrip (montecarlo method). The cluster approach corrects for the multiple comparison problem (Maris and Oostenveld, 2007). Dependent samples t-tests were computed at each sensor and for each cross-frequency interaction. Clusters were obtained by summing up t-values, which were adjacent in space and frequency below a cluster alpha level of 5%. A permutation distribution was computed by randomly switching condition labels within subjects in each of 1000 iterations. The permutation p-value was obtained by comparing the cluster statistic to the random permutation distribution. The observed clusters were considered independently significant when the sum of t-values exceeded 95% of the permutated distribution.

## Results

We applied both coupling measures, namely PAC and AEC to assess theta-gamma and alpha-gamma interactions. In the first study, we found that alpha power was increased during 10 Hz tACS (Helfrich et al., 2014b). Our results indicate that only alpha-gamma PAC was increased during alpha entrainment (t_13_ = 3.77, p = 0.002, d = 1.13; Figure 2), while neither theta-gamma PAC (t_13_ = -1.74, p = 0.11; d = -0.71) nor AECs (theta-gamma: t_13_ = -0.66, p = 0.52, d = 0.04; alpha-gamma: t_13_ = 1.46, p = 0.17, d = 0.61) differed significantly between sham and stimulation. This effect was confined to two parieto-occipital clusters (cluster 1: p = 0.003; cluster 2: p = 0.005; Figure 2). The results demonstrated a regional-specific increase in alpha-gamma PAC during alpha entrainment. To quantify the impact on gamma power, we first analyzed power differences between sham and stimulation and did not observe a significant difference (t_13_ = -0.15, p = 0.88, d = -0.06; Figure 3A). Secondly, we assessed whether the distribution of gamma-power relative to low frequency phase was modulated, i.e. whether gamma power was now preferentially locked to the trough of the low frequency phase (so called nesting; for an illustration see Figure 1B). Hence, we analyzed a power ratio, where alpha-trough-locked gamma power was divided by alpha-peak-locked gamma power and found a significant cluster in parieto-occipital cortex (p = 0.02; d = 0.73; Figure 3B), corresponding to cluster 1 of increased PAC during stimulation (Figure 2). This finding was consistent with an increased nesting of gamma power. We only found one significant cluster, most likely due to the small size of cluster 2, which consisted only of 3 electrodes (Figure 2). This result indicates a frequency-specific locking of gamma power to the trough of the alpha oscillation, since the gamma-power ratio was not modulated relative to the theta-phase (cluster test: p > 0.05, d = - 0.26; Figure 3B).

**Figure 2:**
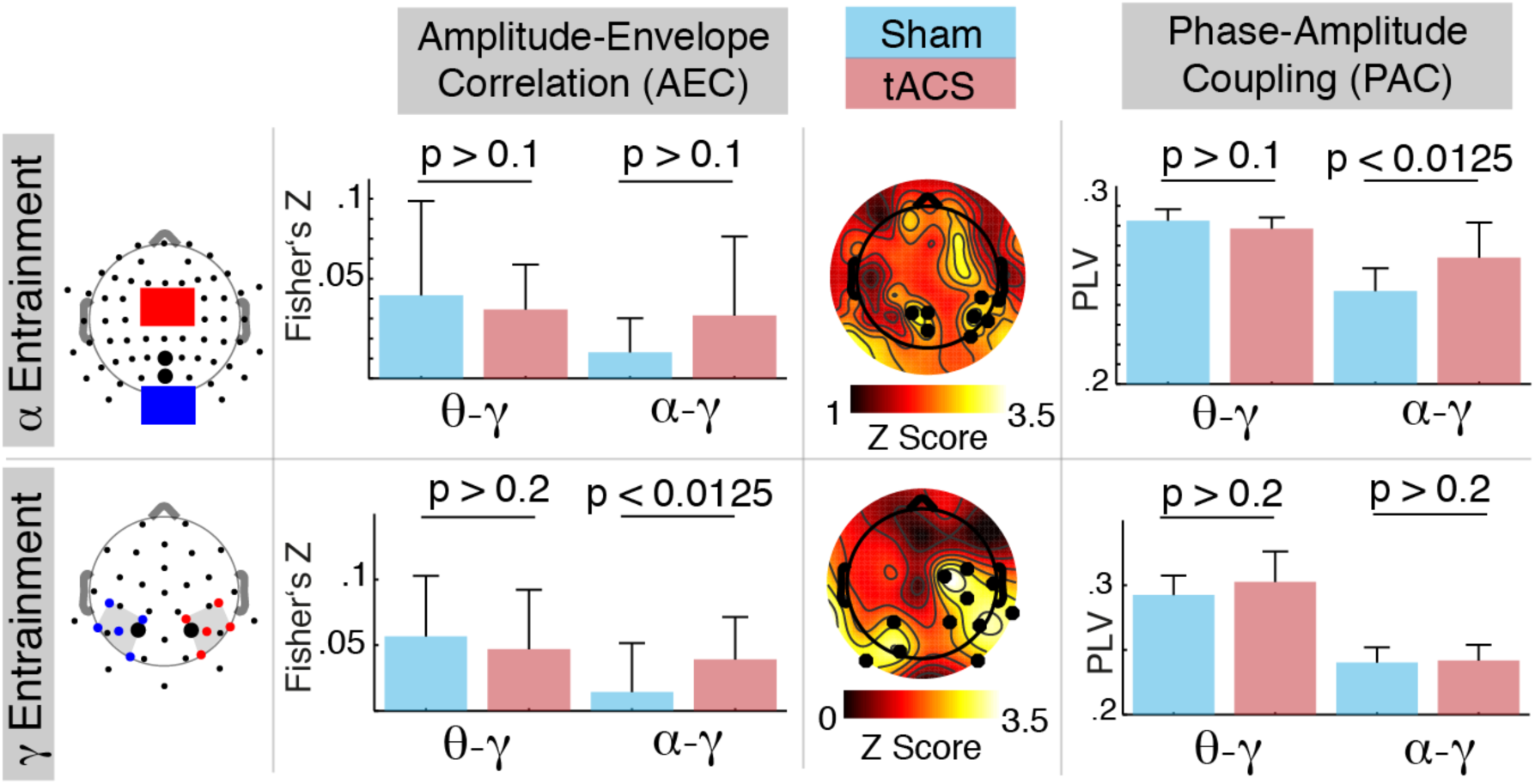
Intrinsic coupling modes of cross-frequency interactions. Upper row: Depicts the electrode montage of study 1 (Helfrich et al., 2014b; electrodes-of-interest are bold; red and blue electrodes depict tACS electrodes), AEC and PAC (mean ± STD) results along with the topography highlighting two significant clusters (black dots) of increased alpha-gamma PAC during 10 Hz tACS. Lower row: Electrode montage, AEC and PAC results from a study 2 (Helfrich et al., 2014a). The topography depicts the increased alpha-gamma AEC during 40 Hz tACS.

**Figure 3:**
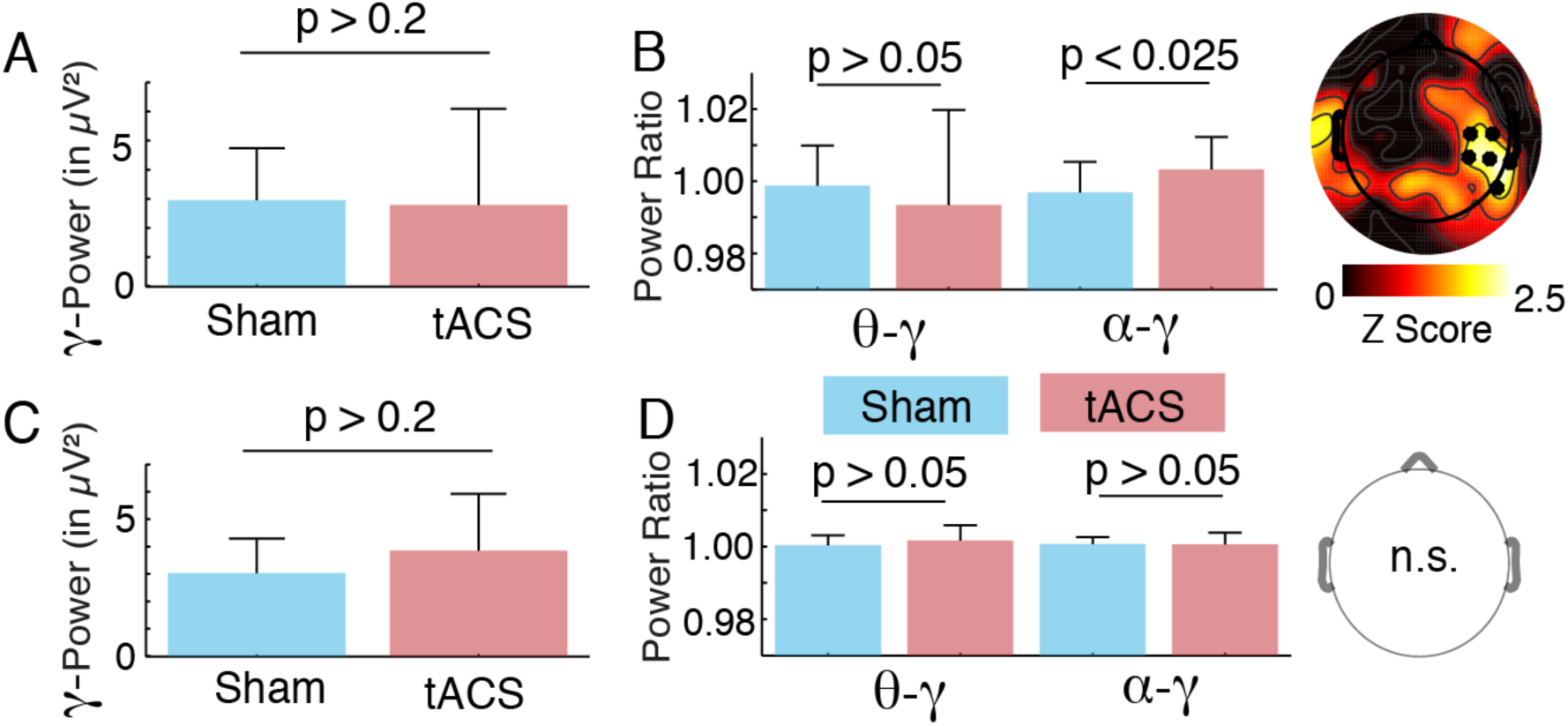
Spectral power and nesting results. (A) The grand-average gamma power results from study 1 (Helfrich et al., 2014b; electrodes of interest) indicate that there was no significant gamma power difference between sham and stimulation. (B) One significant cluster of increased gamma power nesting in the alpha phase during 10 Hz tACS in study 1 was found over lateral parieto-occipital sensors, corresponding to cluster 1 of increased PAC during stimulation (Figure 2). (C) Grand-average gamma power results from study 2 (Helfrich et al., 2014a; electrodes of interest) indicate that gamma power did not differ between sham and stimulation. (D) No gamma power nesting effects were observed during 40 Hz tACS.

In the second study, gamma oscillations were selectively entrained by 40 Hz tACS without concomitant gamma power changes (t_13_ = 1.23, p = 0.24, d = 0.47; Figure 3C). However, a prominent alpha power decrease was observed in states of exogenously entrained gamma-band signatures through enhanced AEC (Helfrich et al., 2014a). Here, we replicated this key finding (alpha-gamma AEC: t_13_ = 3.82, p = 0.002, d = 0.71) and confirmed the regional specificity (cluster test: p = 0.005). This effect was also frequency-specific (theta-gamma AEC: t_13_ = -0.73, p = 0.47, d = -0.22) and was not accompanied by changes in PAC (theta-gamma: t_13_ = 1.30, p = 0.22, d = 0.50; alpha-gamma: t_13_ = 0.44, p = 0.67, d = 0.12). Subsequently, no nesting differences were observed between sham and stimulation (Figure 3D; cluster test: all p > 0.05; theta-gamma: d = 0.37, alpha-gamma: d = -0.04).

## Discussion

Our results demonstrate that selective driving of alpha or gamma oscillations differentially modulated their physiological interaction. Entrainment of alpha oscillations increased PAC and promoted a preferential phase-locking of gamma power to the alpha trough, while gamma-band entrainment modulated AECs. These results indicate that both can serve as master and slave through distinct coupling modes, possibly working as opponent processes to subserve different cortical functions (Schroeder and Lakatos, 2009).

Phase or amplitude coupling are regularly observed in electrophysiological data and have recently been conceptualized as intrinsic coupling modes (ICMs; Engel et al., 2013), which possibly subserve different cortical computations. While amplitude coupling might regulate the activation of distributed neuronal populations, phase coupling is thought to mediate the inter-areal cortical information flow (Engel et al., 2001; Fries, 2005; Hipp et al., 2012; Engel et al., 2013). The present results extend this concept and suggest that interactions between both coupling modes might constitute a feedback mechanism that balances the relation between slow and fast oscillatory rhythms and, thus, optimizes the cortical processing dynamics.

These results are in line with previous correlative evidence, which suggested that coupled alpha and gamma oscillations constitute the functional architecture of the visual system and subserve sensory input selection, attentional control or short-term visual memory (Spaak et al., 2012; Jensen et al., 2014). Hence, the present results provide causal evidence for the physiological alpha-gamma interplay (Osipova et al., 2008; Voytek et al., 2010; Jensen et al., 2014), given that all effects were limited to their interaction in the stimulated parieto-occipital cortex during visual tasks.

So far, tACS has been suggested to operate in canonical frequency boundaries via directed entrainment of ongoing oscillatory activity (Herrmann et al., 2013). Recently, several studies suggested that secondary effects might occur across different temporal scales in coupled or nested neuronal networks (Boyle and Fröhlich, 2013; Wach et al., 2013; Helfrich et al., 2014a). The selective up-regulation of alpha activity by alpha-tACS (entrainment; Herrmann et al., 2013; Helfrich et al., 2014b) or the selective down-regulation of alpha activity by gamma-tACS (enslavement; Boyle and Fröhlich, 2013; Helfrich et al., 2014a) will allow to extend previous correlative evidence for the functional role of alpha oscillations (Jensen et al., 2014).

On the one hand, the present results indicate that tACS can be used as a powerful tool to assess the causal role of CFC for cortical computations, in light of the various pitfalls that hamper the correlative analysis of CFC (Aru et al., 2014). On the other hand, it is the nature of non-invasive brain stimulation that its effects are often small (Thut et al., 2011). In addition, these results were obtained from two different studies, in which participants were engaged in different visual tasks (Helfrich et al., 2014a, 2014b), impeding definitive conclusions. However, they provide the first evidence that tACS modulates oscillatory brain activity across several temporal scales and therefore, might extend our understanding of the mechanisms, which orchestrate cross-frequency coupling (Schroeder and Lakatos, 2009; Engel et al., 2013).

Taken together, the results of both studies indicate that distinct coupling modes are causally involved in different cortical computations and that the rich spatiotemporal correlation structure of the brain might constitute the functional architecture for cortical processing and specific multi-site communication.

## Acknowledgements

The authors declare no competing financial interests. This work was supported by grants from the European Union (ERC-2010-AdG-269716, AKE), the German Research Foundation (SFB936/A3/Z1, AKE; SFB/TRR 31, CSH; SPP1665 EN 533/13-1, AKE; SPP1665 HE 3353/8-1, CSH), and the German National Academic Foundation (RFH).

